# SELVa: Simulator of Evolution with Landscape Variation

**DOI:** 10.1101/647834

**Authors:** Elena Nabieva, Georgii A. Bazykin

**Affiliations:** Skolkovo Institute of Science and Technology, Moscow, Russia; Kharkevich Institute of Information Transmission Problems, Moscow, Russia

## Abstract

Organisms evolve to increase their fitness, a process that may be described as climbing the fitness landscape. However, the fitness landscape of an individual site, i.e., the vector of fitness values corresponding to different variants at this site, can change with time due to changes in the environment or substitutions at other epistatically interacting sites. We present SELVa, the Simulator of Evolution with Landscape Variation, aimed at modeling the substitution process under a changing single position fitness landscape in a set of evolving lineages forming a phylogeny of arbitrary shape. Written in Java and distributed as an executable jar file, SELVa provides a flexible framework that allows the user to choose from a number of implemented rules governing landscape change.

**Availability:** https://github.com/bazykinlab/SELVa

## Introduction

The differences between species arise in the course of evolution due to substitutions – mutations that spread to fixation in evolving lineages. The properties of the substitution process are shaped by mutation giving rise to new variants and by selection favoring some variants over others. Computer simulation methods have established themselves as invaluable tools to infer the characteristics of these processes (for review, see (Arenas, 2012); for a catalogue see (Peng et al., 2013)). Over the years, simulation methods have grown increasingly sophisticated in the models that they implement: from evolving nucleotide, codon or amino acid sequences only by substitutions governed by one of several well-established models (Yang, 1997; Rambaut and Grass, 1997) to allowing insertions and deletions (Fletcher and Yang, 2009; Strope et al., 2007), to modeling evolution under domain preservation or structural constraints (Koestler et al., 2012); and further on. These programs generally permit the user to model inhomogeneity of the evolutionary process among different sites; they may do so, for example, by designating certain sites or groups of sites as invariable or, conversely, as having increased or decreased rates of evolution.

An important feature of evolution is that its parameters themselves change with time and/or along different phylogenetic branches. Fitness landscapes change over time, whether reflecting environmental changes or, in the case of single-position fitness landscapes, epistatic interactions with other sites that are also undergoing change (Bazykin, 2015). For example, a study of evolution in *Drosophila* found that fitness fluctuates at rates comparable to those of nucleotide changes (Mustonen and Lässig, 2007). Studying the effect of changing fitness landscapes therefore calls for an evolutionary simulator capable of modeling landscape changes. While a recently published individual-based forward simulator of population dynamics, SANTA-SIM (Jariani et al., 2019) accommodates a number of scenarios for changing selection, in the realm of sequence evolution simulation along phylogenies, this niche is, to the best of our knowledge, yet to be filled. Some existing simulators do allow model parameters to vary among branches (Fletcher and Yang, 2009; Hall, 2008; Strope et al., 2007; Yang, 1997, p. 199) or at a specified time on a branch (Sipos et al., 2011), yet these capabilities are rather limited. We therefore developed SELVa, the Simulator of Evolution with Landscape Variation. What distinguishes SELVa from existing evolutionary simulators is its focus on the effects of landscape change. This is achieved by its ability to flexibly parametrize landscape change, giving the user multiple options to specify when, where and how the landscape change occurs.

## Results

### Implementation and usage

SELVa is implemented in Java and distributed as an executable jar file. It requires installation of no additional packages.

SELVa simulates the molecular evolution as the process of substitution in a string of a specified length, with positions corresponding to nucleotides, amino acids, or indeed any type of sequence variants. The user provides the program with a rooted phylogenetic tree in Newick format and a configuration file detailing the options of the simulation. Positions are split into one or more classes, each characterized by the same vector of fitness values of individual variants that can change in the course of simulation. The simulation starts with an initial fitness landscape encoded as vector(s) and either supplied by the user in a file or generated from one of the supported probability distributions (currently, they include the gamma and the lognormal). At the start of the simulation, SELVa generates the root state for each position by sampling from the stationary distribution corresponding to the initial fitness vector (Yang 2006, Yang and Nielsen 2008).

Upon completion, SELVa prints out the sequence(s) generated at each node of the phylogenetic tree in the course of the simulation, and, optionally, the times of landscape changes and the fitness vectors generated at those times, providing the user with detailed information about the course of the simulation.

### Landscape change timing

SELVa allows for several regimes governing the timing of landscape changes. They can occur stochastically as a Poisson process (Figure 1a), or deterministically, either at evenly spaced time intervals (Figure 1b and b’) or at user-specified positions in the tree (Fig 1c). In the deterministically evenly-spaced case, the landscape values can be independent between branches (1b) or shared by parallel branches (1b’).

**Figure 1.**
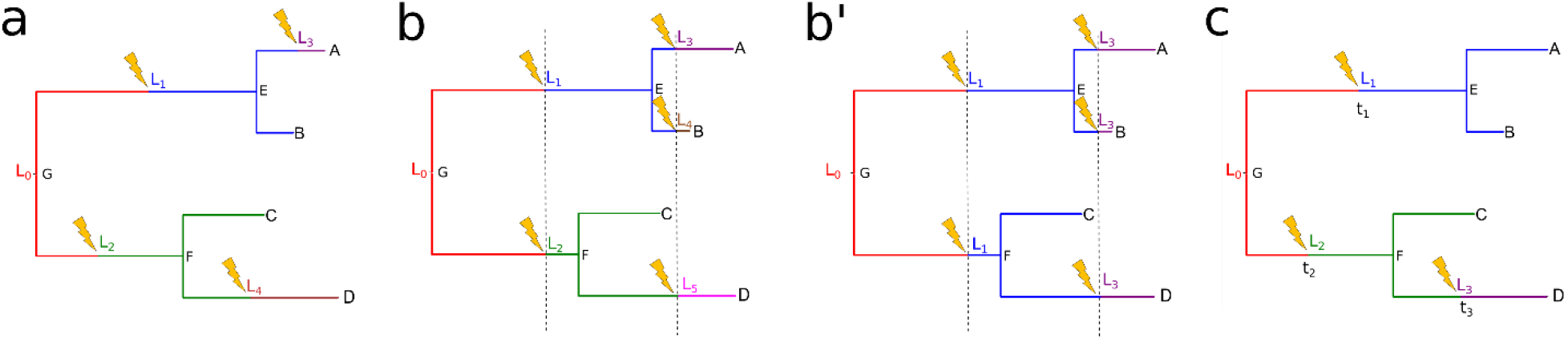
Landscape change options. The landscape change regimes currently supported by SELVa. “Lightning strikes” denote landscape change evens; colors and L_i_ labels correspond to landscapes. L_0_ is the initial landscape in all scenarios. a. The landscape change occurs stochastically. b and b’. The landscape change occurs at evenly spaced time intervals, and the values of the landscapes are independent for different branches (b) or shared among parallel branches (b’). c. The landscape change occurs at user-specified times t_1_, t_2_, t_3_.

### Generation of the new landscapes

The values of the new fitness vector can be independently sampled from the same distribution as the initial fitness vector; they can be obtained from the previous fitness vector by randomly permuting its values; they can be explicitly specified by the user (currently, only if the landscape change times are also manually specified by the user); or they can be derived from the previous fitness vector by either increasing or decreasing the fitness of the current allele at the site. The last option permits the modeling of processes in which the fitness of the current allele increases or decreases with time, e.g., as a result of epistatic interactions with other sites. Other rules for selecting the new landscape are in the planning.

### Parallel simulations

The user has the option of carrying out multiple parallel independent simulations under the same settings in a single execution. If multiple processors are available, these parallel simulations can be divided among multiple threads.

The full options and their details are given in the Manual (available in the github repository).

## Conclusion

SELVa is a pioneering evolutionary simulator focused specifically on modeling the effect of changes in the fitness landscapes. By providing the user with a variety of options for specifying the regime and the specifics of fitness landscape changes, it fills a hitherto vacant need for exploring the effect of single-position fitness landscapes dynamics on the substitution process.

## Acknowledgements

The authors thank Anastasia Stolyarova, Galina Klink, Anfisa Popova, and Dr. Alexey Neverov for being the first SELVa users and providing helpful feedback and suggestions.

We thank the Skoltech Biomedical Initiative for funding.

