## Supplementary materials for "SELVa: Simulator of Evolution with Landscape Variation"

***Simulation details***

SELVa simulates evolution as an event-driven process, following the general simulation framework of Chapter 12.6.1.3 of (Yang, 2014). In particular, a fitness vector is used to derive the substitution rate matrix $Q=\{q_{ij}\}$ that contains the instantaneous rates of substitution from allele $i$ to allele $j$, and the stationary distribution vector π for the current landscape (Yang and Nielsen, 2008). Specifically, the fitness vector is first converted to the unnormalized substitution rate matrix $Q^{raw}$, with $q_{ij,i \neq j}^{raw} = \frac{F_j - F_i}{1 - e^{F_i - F_j}}$ proportional to the instantaneous rate of substitution of allele $i$ to allele $j$, and $q_i^{raw} = q_{ii}^{raw} = -\sum_{j:j \neq i} q_{ij,i \neq j}^{raw}$ the negative of the total rate of substitution *from* allele $i$. Here, $F_i$ is the fitness of allele *i* obtained from the fitness vector. Then, if the alleles are distributed according to the stationary distribution π associated with $Q^{raw}$, the expected substitution rate for the position is $-\sum_i q_i^{raw} \pi_i$. Dividing $Q^{raw}$ by $-\sum_i q_i \pi_i$ produces $Q^{normalized}$, such that $-\sum_i q_i^{normalized} \pi_i = 1$, i.e., the expected substitution rate is equal to 1, i.e., one substitution is expected to occur per site per unit of evolutionary branch length. Following `evolver` (Yang, 1997), normalization is the default behavior. Alternatively, we can choose to scale $Q^{raw}$ to get the expected substitution rate to be 1 only for the flat fitness vector such as `(1,…,1)` whose corresponding $Q^{raw}$ consists of 1's off the diagonal and $-(|A|-1)$ on the diagonal, where |A| is the alphabet size. $Q^{scaled}$ is then obtained by dividing $Q^{raw}$ by $\sum_{|A|} \frac{1}{|A|}(|A|-1) = (|A|-1)$, $\frac{1}{|A|}$ being is the stationary probability of any allele for the flat fitness vector. If the matrix is left unnormalized, the user has the option of scaling the stochastic landscape change rate to it or not. We expect very few users to forego Q normalization.

The substitution process for a single site currently occupied by allele *i* is modeled as a Markov chain with the expected inter-event time $1/q_i$, where $q_i = \sum_{j \neq i} q_{ij}$ is the total rate of transition away from *i*. For longer sequences, the transition rates for all sites are summed to obtain the rate of a substitution event occurring at *any* site, and then the site where a substitution occurs is chosen at random proportionally to the rate of transition away from the allele occupying it. Once a site occupied by an allele *i* is chosen to be mutated, its new allele *j* is selected proportionally to $q_{ij}$. This process is repeated along each branch, starting at the root of the tree; after a substitution occurs, the waiting time until the next substitution is adjusted to reflect the new allele. If the waiting time until the next event exceeds the remaining branch length, no substitution occurs on the remainder of the branch.

We adapt this widely-used (Dalquen et al., 2012; Fletcher and Yang, 2009; Koestler et al., 2012; Sipos et al., 2011) framework to accommodate stochastically or deterministically scheduled landscape changes. In the stochastic case, we add to the set of possible stochastic events (i.e., substitutions at the sequence sites) a landscape change event governed by the user-specified rate. The calculation of the inter-event waiting time and the choice of the next event are then carried out as described above by treating the landscape change event as an extra "site" whose transition rate set by the user. The landscape change event causes a recalculation of the matrix Q and the vector π, and the subsequent substitution process is governed by their new values.

For deterministic landscape change timing, two processes are simulated at the same time: the stochastic sequence substitutions, carried out as described above, and the deterministically-scheduled landscape change. At each step of the simulation, SELVa generates a waiting time until the next Poisson (i.e., substitution) event. Then, this waiting time is compared to the time remaining until the next deterministic landscape change. If the next event is the stochastic substitution event, then it takes place as described above, and the time remaining until the end of the branch or until the next deterministically-scheduled landscape change is adjusted. If the next event is a deterministically-scheduled landscape change, then the change is carried out, and a new stochastic waiting time for the next substitution event is generated using the new landscape. When a branch splits into two children, the next landscape change is deterministically scheduled on both branches at the time remaining until the next deterministic event.

***Performance***

We evaluated the performance of SELVa on a mitochondrial tree of 3558 species. The height of the tree (longest path from root to leaf) is 0.97, and the sum of branch lengths is 95.02. The initial fitness landscape was provided in a file and was set to be (1, 0.1, 0.1, …, 0.1). The transition rate matrix was normalized, so that one substitution was expected to take place per site per unit time. New fitness vectors were generated by permuting the values previous fitness vector. The following landscape change regimes were tested: no change, stochastic change with rate 1.0, stochastic change with rate 10.0. A single parallel simulation for a sequence of length 1, a single parallel simulation for a sequence of length 1000, and 1000 parallel simulations for sequences of length 1 were tested. To evaluate the effect of keeping track of and printing out landscape change times and intermediate landscape values, we tested the performance with and without that option. We averaged the results over 100 independent runs of SELVa (which, for the 1000 parallel simulations case, means 100 000 different actual simulations).

The simulations were run on a Samsung Notebook 9 with Intel Core i7-6500U 2.50GHz processor and 8 GB total RAM. Multi-simulations runs were not parallelized.

The results are summarized in Supplementary Table 1

*Supplementary Table 1: average running times and memory consumption for stochastic landscape change*

| seq length | # parallel simul. | print landscapes? | no change | | stochastic λ = 1 | | stochastic λ = 10 | |
|---|---|---|---|---|---|---|---|---|
| 1 | 1 | yes | 0.35 | 28.8 | 0.42 | 30.1 | 0.45 | 30.2 |
| | | no | 0.32 | 25.6 | 0.34 | 26.3 | 0.36 | 32.7 |
| 1000 | 1 | yes | 1.25 | 42.9 | 1.6 | 43.6 | 1.37 | 52.2 |
| | | no | 1.24 | 43 | 1.3 | 41.9 | 1.27 | 44.7 |
| 1 | 1000 | yes | 9.5 | 776.7 | 14.9 | 840.7 | 28.33 | 1070.1 |
| | | no | 1.21 | 40.4 | 9.3 | 745 | 18.2 | 889.8 |
| | | | **time (sec)** | **memory (MB)** | **time (sec)** | **memory (MB)** | **time (sec)** | **memory (MB)** |

The infrastructure associated with each simulation run is the most computationally expensive aspect of the program, with the infrastructure for keeping track of the landscape information adding some cost to it.
